# A modular approach to handle *in-vivo* drift correction for high-density extracellular recordings

**DOI:** 10.1101/2023.06.29.546882

**Authors:** Samuel Garcia, Charlie Windolf, Julien Boussard, Benjamin Dichter, Alessio P. Buccino, Pierre Yger

**Affiliations:** Centre de Recherche en Neuroscience de Lyon, CNRS, Lyon, France; Columbia University, New York City, NY, USA; CatalystNeuro, Benicia, CA, USA; Allen Institute for Neural Dynamics, Seattle, WA, USA; Institut de la Vision, Sorbonne Université, INSERM, Paris; Lille Neurosciences & Cognition (lilNCog) – U1172 (INSERM, Lille), Univ Lille, CHU Lille 59045 Lille, France

**Keywords:** spike sorting, drift, benchmark, ground-truth, electrophysiology

## Abstract

High-density neural devices are now offering the possibility to record from neuronal populations *in-vivo* at unprecedented scale. However, the mechanical drifts often observed in these recordings are currently a major issue for “spike sorting”, an essential analysis step to identify the activity of single neurons from extracellular signals. Although several strategies have been proposed to compensate for such drifts, the lack of proper benchmarks makes it hard to assess the quality and effectiveness of motion correction. In this paper, we present an exhaustive benchmark study to precisely and quantitatively evaluate the performance of several state-of-the-art motion correction algorithms introduced in literature. Using simulated recordings with induced drifts, we dissect the origins of the errors performed while applying motion-correction algorithm as a preprocessing step in the spike sorting pipeline. We show how important it is to properly estimate the positions of the neurons from extracellular traces in order to correctly estimate the probe motion, compare several interpolation procedures, and highlight what are the current limits for motion correction approaches.

**Significance statement:** 

## Introduction

Recording from increasingly larger neuronal populations is a crucial challenge to unravel how information is processed by the brain. The recent development of CMOS-based high-density multi-electrode arrays (HD-MEAs), both for *in-vitro* [2, 11] and *in-vivo* applications, such as Neuropixels probes [1, 14, 25], has dramatically boosted the yield of extracellular electrophysiology experiments. To maximize the return from these novel probes, the “spike sorting” step, i.e., the process which extracts single-neuron activities from the recorded traces, has also undergone major advances both in algorithmic development [5, 8, 15, 16, 18, 20, 21, 25, 30] and quantitative benchmarks [6, 12, 18].

The layout of high-density probes for *in-vivo* applications usually consists of a long shank (e.g., ∼1 mm for Neuropixels 1.0 [14]) to allow the electrodes to traverse multiple regions of the brain and reach deep structures. Due to the different mechanical properties of the probe and the brain tissue, it is very common to observe a relative movement of the tissue with respect to the probe. This phenomenon is known as *drift*. The origins and types of such drifts can be diverse. For example, cells are likely to slowly drift from initial positions because of the pressure release in the tissue after an acute probe insertion; abrupt and discontinuous drift events could be caused by sudden rig instabilities and movement artifacts. When a neuron moves with respect to the recording electrodes, its waveforms are distorted (Figure 1A), challenging the operation of spike sorting algorithms which mainly rely on waveform similarities to cluster different neurons in the recordings.

**Figure 1.**
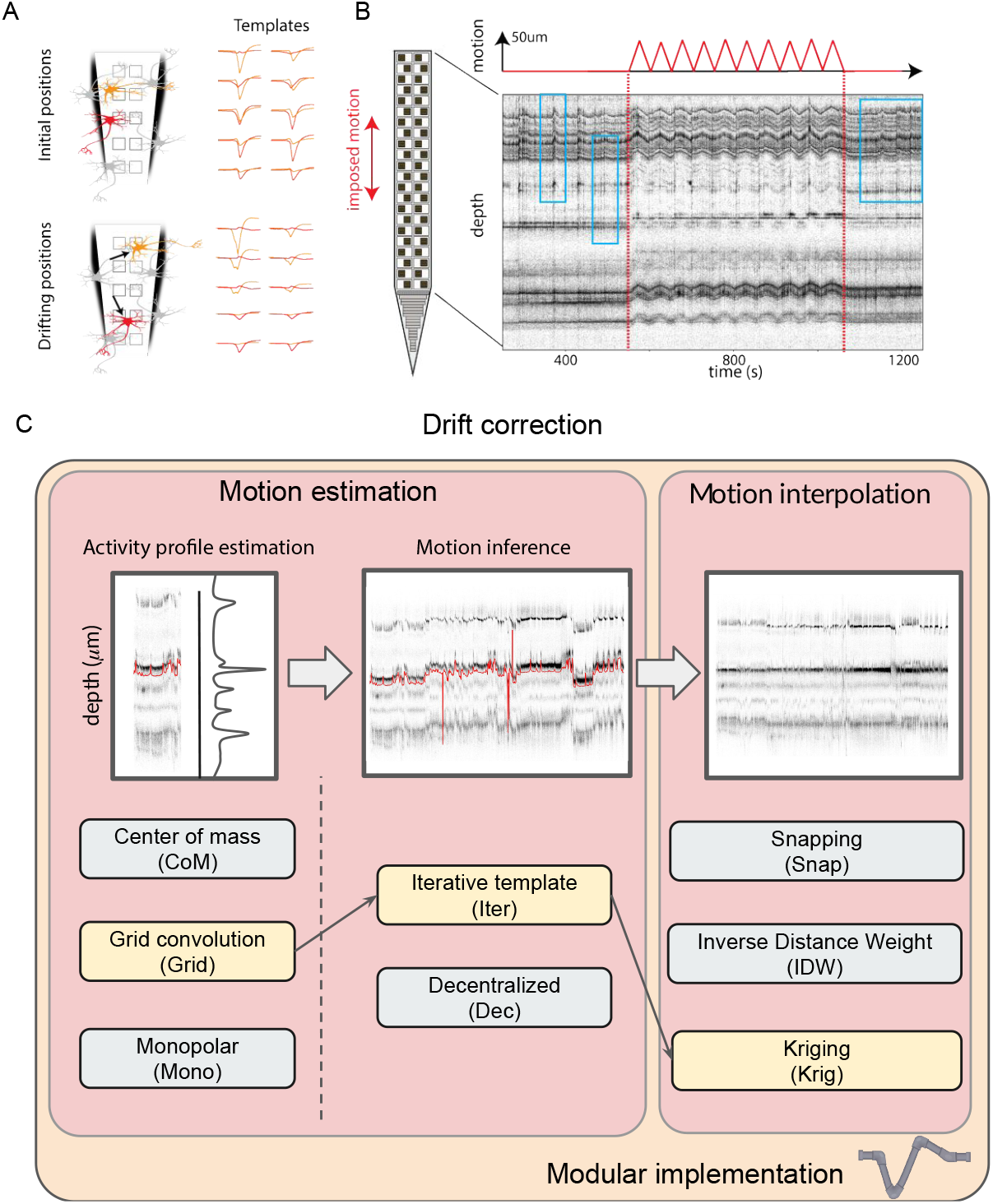
Common pipeline for drift-correction algorithms. **A)** Illustration of the problems raised by neuronal drifts. Two neurons (red and orange) have two distinct waveforms, but after some non rigid drifts (movements), the templates are changed. **B)** Illustration of the drift *in-vivo*. A Neuropixels 1.0 probe in the cortex of the mice is moved up and down with an imposed triangular movement of amplitude 50 μm (top trace) [24, 25] – `p1` dataset. The row shows the depth of all the peaks detected via signal threshold (*positional* raster plot, see text) as function of the time. As one can see, when the probe starts moving (red dotted line), so does the depth of the peaks, since neurons are drifting across channels. While the drift here is imposed, spontaneous drifts can happen locally and non homogeneously over the whole probe (see inside light blue boxes before and after movements). Panels A and B are adapted from [5], with permission. **C)** Key algorithmic steps for the motion correction pipeline. For each step, the various implementations available in `SpikeInterface` are listed. The yellow boxes connected with arrows correspond to a `Kilosort` 2.5/3-like approach [20].

Due to the spatial regularities of the recordings, which makes drifts relatively coherent in space (Figure 1B shows a *raster map* of a recording with an induced triangular probe movement [24]), a common approach is to mimic drift correction procedures that have been applied in the field of calcium imaging [13]. First, motion is estimated from the spiking activity extracted from the recording (Figure 1C - left). Then, the estimate is used to interpolate or register the recording as a preprocessing step (Figure 1C - right). Different methods have been recently proposed to correct for drifts before spike sorting [3, 20, 25, 27], but their evaluation is mainly qualitative. A quantitative evaluation of the performance and effectiveness of different strategies is still lacking, probably due to the complexity of performing such benchmarks. To benchmark drift correction methods, one would need to know the ground-truth motion that is generating drifts. In addition, these methods are usually integrated into complicated spike sorting pipelines, and this makes it hard to track down what part of the error was really tied to the motion correction step.

In this work, we extensively benchmarked state-of-the-art methods for drift correction to solve the above mentioned challenges. To obtain recordings with ground-truth motion, we used biophysically detailed simulations with drifting neurons. This allowed us to systematically explore a wide range of drifting recordings of varying complexity. We further identified and broke down existing drift correction strategies into a modular architecture that allowed us to properly evaluate the performance of different methods for estimating motion, interpolating the traces, and their overall impact on spike sorting outcomes. All the source code used to generate the synthetic recordings and to reproduce results and figures is available at https://github.com/samuelgarcia/spike_sorting_motion_benchmark_ paper_2023.

## Materials and Methods

### A modular implementation of drift correction

In order to properly evaluate and benchmark the latest *in-vivo* drift correction methods available nowadays in the literature [20, 27], we first identified the individual sub-components of the different approaches and implemented a modular system to be able to substitute different steps. In fact, most of the available drift correction methods acting at the preprocessing level [3, 20] share similar ideas, so that the global algorithmic pipeline can be decomposed and summarized as outlined as follows (Figure 1C):

First, a round of peak detection is performed and the putative peak depths (defined as the position along probe’s insertion direction) for each action potential are estimated (this first step is usually done on a subset of peaks to speed up the computation time). The estimated depths are binned in short time windows (e.g., 2 seconds) to obtain an **activity profile** (Figure 1C - left) as function of depth. It is important to stress that a time windows that is too small might not contain enough spikes to properly estimate the activity profile, while too large time windows might result in *blurred* activity profiles because of the drift within each time window. From the activity profiles varying in time, the **motion inference** step aims at recovering the motion signals (red line in Figure 1C - middle). Finally, one can interpolate the traces with the inverse motion signals to compensate for the estimated motion (**motion interpolation**, Figure 1C - right).

The general pattern of these three processing stages is seen across different spike sorting pipelines, though the details and implementations of the different steps vary from algorithm to algorithm. For example, in [20, 25], an average activity template is computed as the mean of activity histograms computed over all temporal windows and used as a reference for motion registration. Nevertheless, this *average template* strategy assumes a stationary spiking activity over the course of the recording. Recent work have proposed a decentralized method [27, 29] to relax this assumption, which uses all the pairwise displacements between activity profiles, instead of using a single reference template, to directly estimate a motion signal to register the traces.

We used the modular architecture of the `sortingcomponents` module of the `SpikeInterface` package [6] to factorize and benchmark all the steps in a modular manner. As shown in the bottom of Figure 1C, we have implemented three major algorithms to estimate the positions of the neurons (see Methods), two methods to estimate the motion given these positions (the one from `Kilosort`, termed “ìterative template” and the one from [3] termed “decentralized”, see Methods), and three methods to interpolate the traces.

This modular implementation allows us first to benchmark the different drift correction steps in a sorter-agnostic manner. It is worth highlighting that these correction methods can be applied as a preprocessing step before any spike sorting algorithm. In addition, the modular implementation can also replicate existing drift correction strategies, including the one similar to `Kilosort` (since v2.5)^1^ (shown in yellow in Figure 1C), and the motion estimation introduced in [3, 27] (note that in this case the motion interpolation step was not used).

### Motion estimation

As illustrated in Figure 1C, motion estimation includes two distinct steps: activity profile estimation and motion inference.

#### Activity profile estimation (peak localization)

To estimate an histogram of the activity of the cells in a given time window *τ*_*win*_ (default is 2 seconds) as function of depth, we used, in all cases, the estimated peak depths obtained via peak localization. We first detected peaks as negative thresholds crossings with a spatio-temporal exclusion zone (see the “locally-exclusive” method of `SpikeInterface`, with parameters *detection*_*threshold* = 10, *local*_*radius*_*um* = 50 μm and *exclude*_*sweep*_*ms* = 0.2ms). From the detected peaks, we estimated their depths in three ways:

##### Center of Mass (CoM)

Assuming that the spike *i* has its waveform 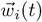 defined on several channels *c* ∈ {1, …, *n*_channels_}, we computed the peak-to-peak values ptp_*i*_(*c*) on every channel. Since every channel has a physical position 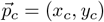 in the 2D space, we can obtain, for every spike *i* its Center of Mass CoM(*i*), as

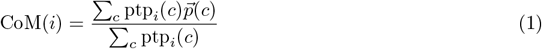

##### Grid convolution (Grid)

Introduced in [20], the idea behind this localization method is to create an exhaustive basis of artificial templates 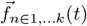 at known positions 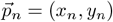, and to estimate the position of a given spike 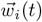 projected onto this basis. If the spatial resolution of this grid is finer than the one of the recording channels, one could expect to enhance the resolution of the localization estimates. First, the scalar products 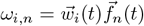 of the spike with all the templates in the basis are computed; then, the spike position Grid(*i*) is estimated as a weighted sum of the positions, with weights equal to the scalar product between the spike waveforms and these templates (negative scalar products are discarded):

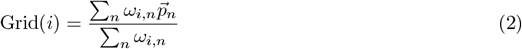

To create the artificial templates, the typical waveform *H*(*t*) of a spike on a single channel is estimated as a median over 1000 normalized waveforms, and then duplicated on all nearby channels, with a spatial decay in amplitude *σ* such that on every channel *c* at a position 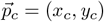 we have:

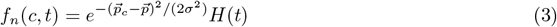

To extend the basis, one can use multiple *σ* values (in the range of 10 to 50 μm). Moreover, in order to reduce the spreading of the scalar products if too many templates are in the basis, we only perform the estimation in Equation 2 on the top 10% of the scalar product values.

##### Monopolar triangulation (Mono)

The idea of this method is to consider the cell as a monopolar current source [3, 7] and infer its position by triangulation given the amplitudes of the templates recorded on all channels. Assuming the spike behaves as a monopolar current source, the extracellular amplitude on a channel *c* generated by a neuron *i* at position 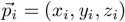 can be expressed as:

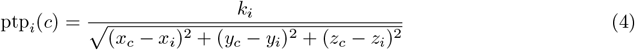

with the *k*_*i*_ term including the magnitude of the current and propagation properties of the tissue (see [7]). Therefore, to estimate the location of the spike one can simply solve an optimization problem with cost function Φ(*x, y, z, k*) and find the (*x*_*i*_, *y*_*i*_, *z*_*i*_) estimated position (and (*k*_*i*_)) from the monopolar approximation by minimization:

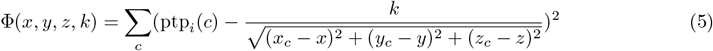

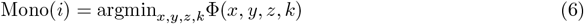

To minimize the cost function, we used the `scipy.optimize` [28] function with the BGFS algorithm.

#### Motion inference

From the activity profiles, the next step is to infer motion. Although the spike localization methods return 2D (CoM, Grid) and even 3D (Mono) locations, the motion inference step currently only uses the *y* coordinate, i.e., the spike depth. For this step, we considered two different methods:

##### Iterative template (Iter)

The first method is the one implemented in `Kilosort` 2.5 [20, 25]. From the activity profiles, a 3D histogram is constructed using the spike counts, spike depths, and log-transformed amplitude as dimensions. Next, a rigid registration is computed by iteratively finding the shift that maximizes the correlation between each bin and a target *average* template (using shifts of *±*30 μm by default), which is also iteratively updated (for the first iteration, an activity profile sample at the center of the recording is used). Next, the depth is divided into sub-blocks to refine the estimation by taking into consideration non-rigid effects. For each block, a similar procedure, but without iteration, is performed to find the best alignment for each spatial window.

##### Decentralized (Dec)

The second method is from [27]. First, it constructs a 2D motion histogram from the activity profile, by binning peak locations over depths. The main difference between this approach and the iterative template one is that the correlation is not computed with respect to a target template, but, instead, a correlation value is computed between each pair of bins (optionally, a time *horizon* can be used to limit the correlation computations to bins within a certain interval to speed up the computation, e.g. 120 s). This results in a pairwise correlation matrix, which is then used to estimate the motion that maximizes all pairwise correlations. To achieve non-rigid motion estimation, the same approach is applied on spatial windows and a spatial prior is used to enforce smoothness between consecutive spatial blocks [29].

### Motion interpolation

Once the drift vectors have been estimated, the next step of the pipeline is to interpolate the extra-cellular traces. Given all the values of the signals *s*_*c*_(*t*) at channels *c* at positions 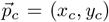, the goal is to determine the value *s*(*t*) of the extracellular traces at a drift corrected position 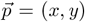 which accounts for the estimated motion. Here again, several methods to interpolate the extracellular traces from the motion vector have been implemented. To be more efficient, interpolation kernels are computed at the temporal resolution of the time histograms used to compute the activity profiles in a chunk-wise manner, not at every time point.

#### Snapping (Snap)

In this case, given an estimated position, the value of the the extracellular traces at the closest electrode from the drift vector is used:

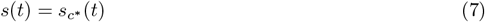

with *c*\* such that we have 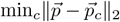.

#### Inverse Distance Weighting (IDW)

The interpolated value is a weighted sum of the values in the vicinity, with weights determined by the distances:

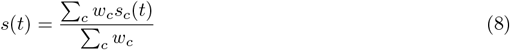

with 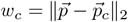. In practice, this weighed sum is restricted to the 3 nearest channels.

#### Kriging (Krig)

Finally, we also implemented the kriging method, as used in `Kilosort` [20]. To be more precise, we computed *K*_*xx*_ as the exponentiated negative distance matrix between all the positions of the known channels 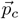. To account for the different x a nd y s cales o f t he Neuropixels layout, `Kilosort` uses different s caling f actors *σ* _*x*_ a nd *σ* _*y*_, a nd w e r euse t he e xact s ame f ormula for comparison purposes. However, a more generic framework has been implemented in `SpikeInterface` to account for the general distances. The distance considered for *K*_*xx*_ in this paper is thus a city-block distance between all channels *i, j* one such that:

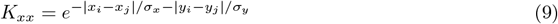

In practice, we set *σ*_*x*_ = 20 μm and *σ*_*y*_ = 30 μm. Then we computed *K*_*yx*_ as exponentiated negative the distance matrix between 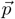 and all the channels 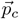. The kriging kernel *W* is obtained as:

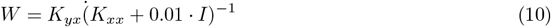

The kernel is sparsified so that all values below ϵ = 1*e*^−3^ are set to 0 and normalized so that each column of *W* sums up to 1.

### Simulated datasets

We used the `MEArec` simulator [4] (version 1.9.0) to generate 10-minute-long synthetic ground-truth recordings. `MEArec` uses biophysically detailed multicompartment models to simulate extracellular action potentials, or so-called “templates”. For this study, we used 13 *default* cell models from layer 5 of a juvenile rat somatosensory cortex [19, 23] to simulate a dictionary of biologically plausible drifting templates on a Neuropixels probe layout (cropped to 128 electrodes in four columns and hexagonal arrangement, a x- and y-pitch of 18 μm and 22 μm, respectively, and an electrode size of 12 μm per side). The drifting templates (100 for each cell model) were obtained by moving the cell models on along a 100 μm straight vertical line with 1 μm steps starting from random initial positions. For each recording, we then selected 256 neurons and generated corresponding spike trains. Templates and spike trains were then convolved following the drift signals and adding a slight modulation in amplitude to add physiological variability. Finally, uncorrelated Gaussian noise with 5 μV standard deviation was added to the traces. The sampling rate of the simulated recordings was set to 32 kHz.

To challenge the drift correction algorithms, we generated various drift recordings using different drift signals, depth distributions, and firing rate profiles.

#### Drift signals

Different types of drift might challenge the drift correction approaches in different ways. We therefore generated three drift signals:

- **Zigzag (Rigid):** For this drift mode we used a rigid drift (all cells move homogeneously across depth) with a triangular slow oscillation, an amplitude of 30 μm and a 30 μm/*min* speed. Figure 2A-B show sample raster maps using this drift mode.
- **Zigzag (Non-rigid):** This mode is similar to Zigzag (Rigid), but in this case the drift is non-rigid, leading to a non-homogeneous movement of all the cells as function of their depths. In this non-rigid case, the drift vector is modulated by a linear gradient, ranging from 0.4 (12 μm maximum displacement) at the upper part of the probe to 1 (30 μm maximum displacement) at the bottom. Figure 2C-D show sample raster maps using this drift mode.
- **Bumps (Non-rigid):** To reproduce the abrupt transient drifts that can often be observed *invivo* [9], we also modelled fast, abrupt drifts that can happen irregularly during the recording. These abrupt events happened at random times (between 30 and 90 s) and moved all cells non-rigidly between -20 and 20 μm at the top and -40 and 40 μm at the bottom of the probe. The drift signals also experience an additional small 3-μm sinusoidal modulation with 40 Hz frequency. Figure 2E-F show sample raster maps using this drift mode.

**Figure 2.**
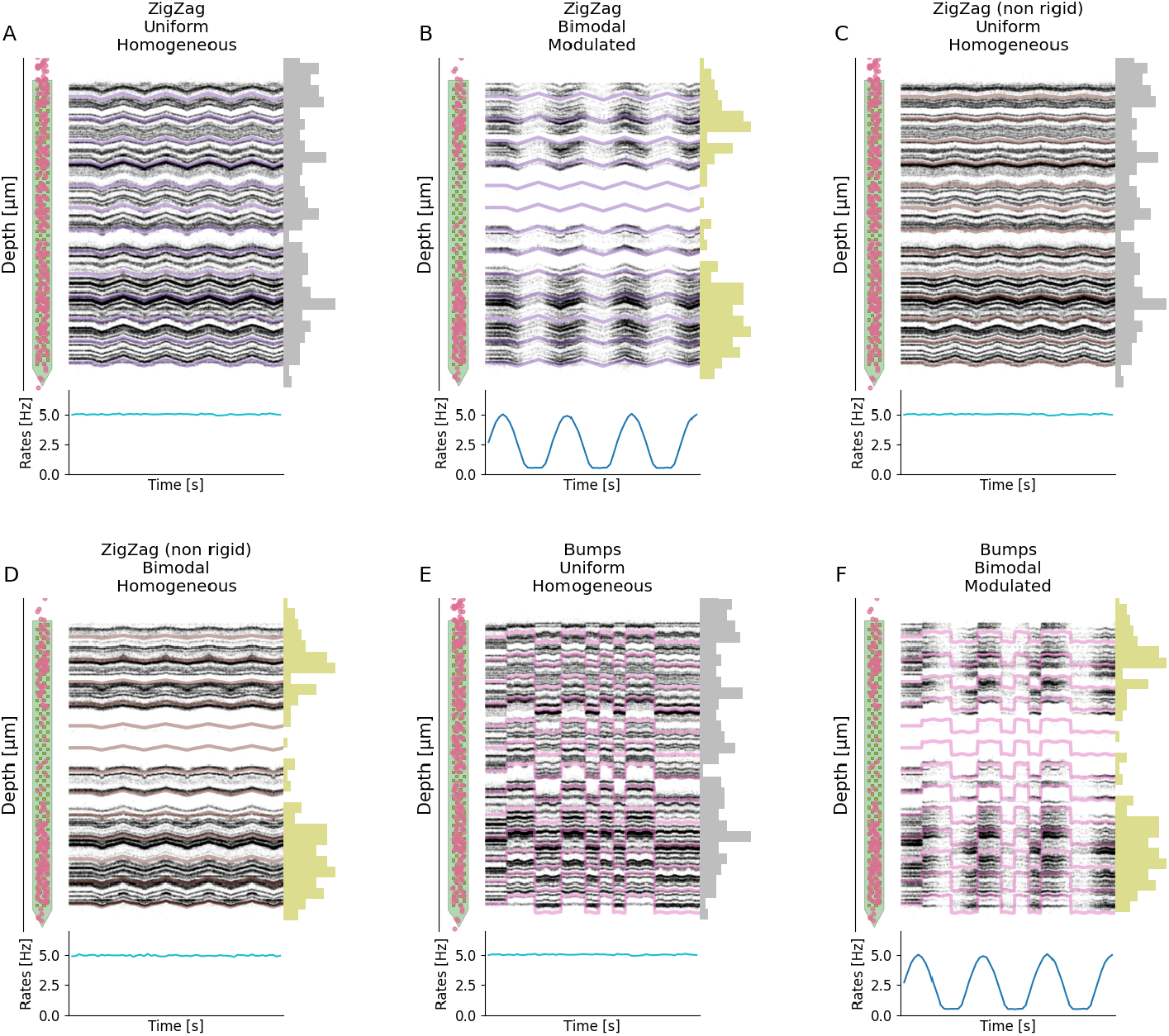
Examples of simulated drift recordings. For each panel, the top shows the layout of the probe with superimposed starting position of each cell (left), the positional raster plot with overlaid ground-truth motion signals (center), and the depth distribution of the 256 neurons. The bottom part displays the firing rate modulation. Here we display 6 out of 12 recordings (the remaining 6 can be found in S1). **A)** Rigid ZigZag drift with uniform depth distribution and homogeneous firing rates. **B)** Rigid ZigZag drift with bimodal depth distribution and modulated firing rates. **C)** Non-rigid ZigZag with uniform depth distribution and homogeneous firing rates. **D)** Non-rigid ZigZag with bimodal depth distribution and homogeneous firing rates. **E)** Bumps with uniform depth distribution and homogeneous firing rates. **F)** Bumps with bimodal depth distribution and modulated firing rates.

#### Depth distributions

The distribution of neurons along the probe can potentially challenge drift correction strategies, as they might result in portions of the probe experiencing lower spiking activity. This effect mimics the uneven distribution of neurons across the brain (e.g., in the cortex).

- **Uniform:** The depths of the selected neurons (256) were drawn from a uniform distribution, as in Figure 2A, C, and E.
- **Bimodal:** The neuron depths follow a bimodal distribution with two peaks at the top and bottom of the probe and a central region with a lower density of cells, as in Figure 2B, D, and F.

#### Firing rates

Drift correction methods may assume stationary firing rates. We therefore simulated two scenarios:

- **Homogeneous:** The activity of all neurons were modeled as homogeneous Poisson sources with 5 Hz mean firing rate, as in Figure 2A, C, D, and E.
- **Sine-modulated:** The spike trains were obtained following a nonhomogeneous Poisson distribution modulated with a slow sine wave with a 3-minute period, as in Figure 2B and F.

We generated a total of 12 scenarios by combining the above mentioned option (3 drift signals *x* 2 depth distributions *x* 2 firing rates). Six of these cases are shown in Figure 2, the rest can be found in the Supplementary Figure S1. In all cases, the drift started after 60 s, so that the first portion of the recording is drift-free. Owing to the reproducibility option of the `MEArec` simulator, which allows one to set random seeds to control all steps of the simulation, for each scenario we generated a *static* counterpart, with the exact same neurons, spiking activity, and noise profile, but only without drifts. We used these *static* recordings to benchmark the motion interpolation step.

### Evaluation

To evaluate the impact of the motion estimation step, we took advantage of the known ground-truth motion signals at different depths. We computed the estimation error as the Euclidean distance between the ground-truth motion and the one estimated by the different methods. Note that to compensate the offsets between estimated and ground truth motions, we aligned the medians of the ground truth and the estimated motions. From these error signals we could then visualize both the evolution of the error over time and the error profile along the depth dimension.

To benchmark motion interpolation, we used the ground-truth motion signals to interpolate the traces and then utilized the ground-truth spiking activity to extract single spike waveforms from the motion-corrected and associated *static* recording. Assuming that the neuron *i* has fired *N* spikes with waveforms 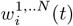 (defined on a subset of channels), we calculated the dispersion around the mean template *T*_*i*_(*t*) as the standard deviation *σ*_*i*_, to measure of how “variable” these spikes are. Normalization is achieved by dividing *σ*_*i*_ by the root mean square of the template *T*_*i*_. To compare the effect of different motion interpolation procedures, we further computed the ratio 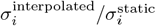 for every neuron *i*. The rationale for such a metric is that spike sorting algorithms assume that waveforms are “sterotypical”, but drift is increasing the variability due to movements. Interpolation should therefore reduce this extra variability introduced by drifts.

Finally, and most importantly, we evaluated how drift correction affects spike sorting outcomes. To do so, we focused on the `Kilosort` 2.5 algorithm, which first introduced drift correction as a preprocessing step to spike sorting [25]. We ran all spike sorting jobs in the `SpikeInterface` framework [6] on a 40-core Intel(R) Xeon(R) Silver 4210 CPU @ 2.20GHz, with 64GB of RAM and an 8GB Quadro RTX 4000 GPU. We used default parameters, but ran the preprocessing and motion correction steps in SpikeInterface instead of `Kilosort` 2.5. To match the `Kilosort` 2.5 processing, before motion correction, we applied a highpass filter (cut-off frequency at 150 Hz), a common median reference (CMR), and finally a local whitening step [20, 30] (using, for each channel, the neighbor electrodes within a 150-μm radius).

Knowing the ground-truth spiking activity, we could compute the *accuracy* of each ground-truth unit *i* as:

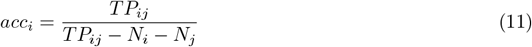

where *j* is the sorted unit matched to the Ground Truth (GT) unit *i, N*_*i*_ and *N*_*j*_ are the number of spikes in the GT and matched sorted unit, respectively, and *T P*_*ij*_ is the number of *true positive* spikes, i.e., the spikes found both in the GT and sorted spike trains. From this accuracy metric, we further classified spike sorted units as:

- *well detected* : units with an accuracy greater or equal than 80%.
- *overmerged* : units with an agreement above 20% with more than one GT unit.
- *redundant* : units with an agreement above 20% with a GT unit that are not the best matching unit. These units can be either oversplit or duplicated sorted units.
- *false positive*: sorted units with an agreement below 20%.
- *bad* : the sum of *overmerged, redundant*, and *false positive* units.

## Results

### Benchmarking motion estimation

To benchmark the performance of different motion estimation strategies, we generated several artificial recordings with 256 neurons and various drift cases (see Simulated datasets). In Figure 3, we first show the results for a representative subset of six drift recordings (one per row). All the remaining cases can be found in Supplementary Figure S2. The left column displays the raster maps with the overlaid drift signals at multiple depths, which give a qualitative view of the drift over the entire recording. The right part of the figure focuses on the errors of different *peak localization* + *motion inference* combinations: the left panels show the errors over time, the central panels the error distributions, and the right panels the errors over probe depth (left - bottom, right - top). The first thing that we should highlight is that motion estimation, irregardless of the method, is generally good. Average errors for relatively smooth drifts (A to D) are less than 5 μm, which is well below the inter-electrode distance. Even in the case of *Bumps* (E-F), where errors are more pronounced, some but not all methods can achieve average errors below 5 μm. Second, the errors tends to be larger at the borders of the probe. This is expected, because we have less spatial information there, thus the motion inference is less accurate. Another general observation is related to the peak localization methods, used to estimate the activity profile: the Center of Mass (*CoM* - blue colors) method yields higher errors, and hence performs worse, than all other estimation methods (but it is the fastest – see figure 4C). The monopolar triangulation (*Mono* - oranges) and the grid convolution (*Grid* - greens) perform similarly, with the former achieving slightly lower error distributions. As for the motion inference step, the decentralized method (*Dec* - dark shades) generally produces lower errors than the iterative template (*Iter* - light shades) approach, except for the ZigZag/Bimodal/Modulated case (panel B). We should also note that the errors from the decentralized method, despite lower on average, show increased variance, due to some time bins being highly misestimated (see *spikes* in the second column of Figure 3B, E, F). During these particular time bins, the increased errors can dramatically affect the motion interpolation step and the overall spike sorting results.

**Figure 3.**
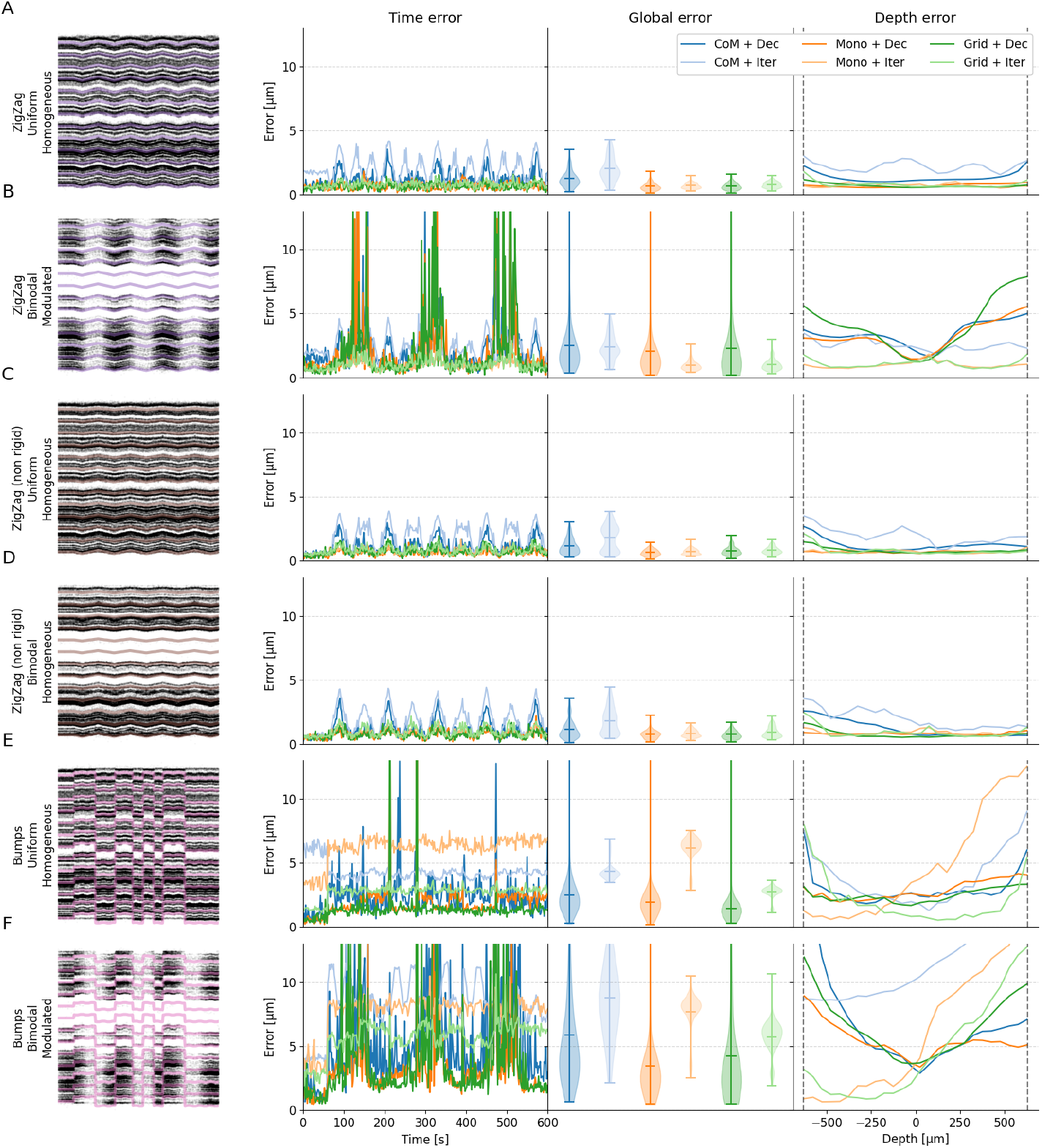
Performance in motion estimation for various drifts scenarios. For various situations of the drifts with overlaid ground-truth motion (left column), errors (from left to right) as function of time (time error), averaged (global error), or as function of the depths (depth error) and for various motion estimation pipelines. Here we display 6 out of 12 recordings (the remaining 6 can be found in Figure S2). **A)** Zigzag (rigid), uniform positions, homogeneous rates **B)** Zigzag (rigid), bimodal positions, modulated rates **C)** Zigzag (non-rigid), uniform positions, homogeneous rates **D)** Zigzag (non-rigid), bimodal positions, homogeneous rates **E)** Bump (non-rigid), uniform positions, homogeneous rates **F)** Bump (non-rigid), bimodal positions, modulated rates

**Figure 4.**
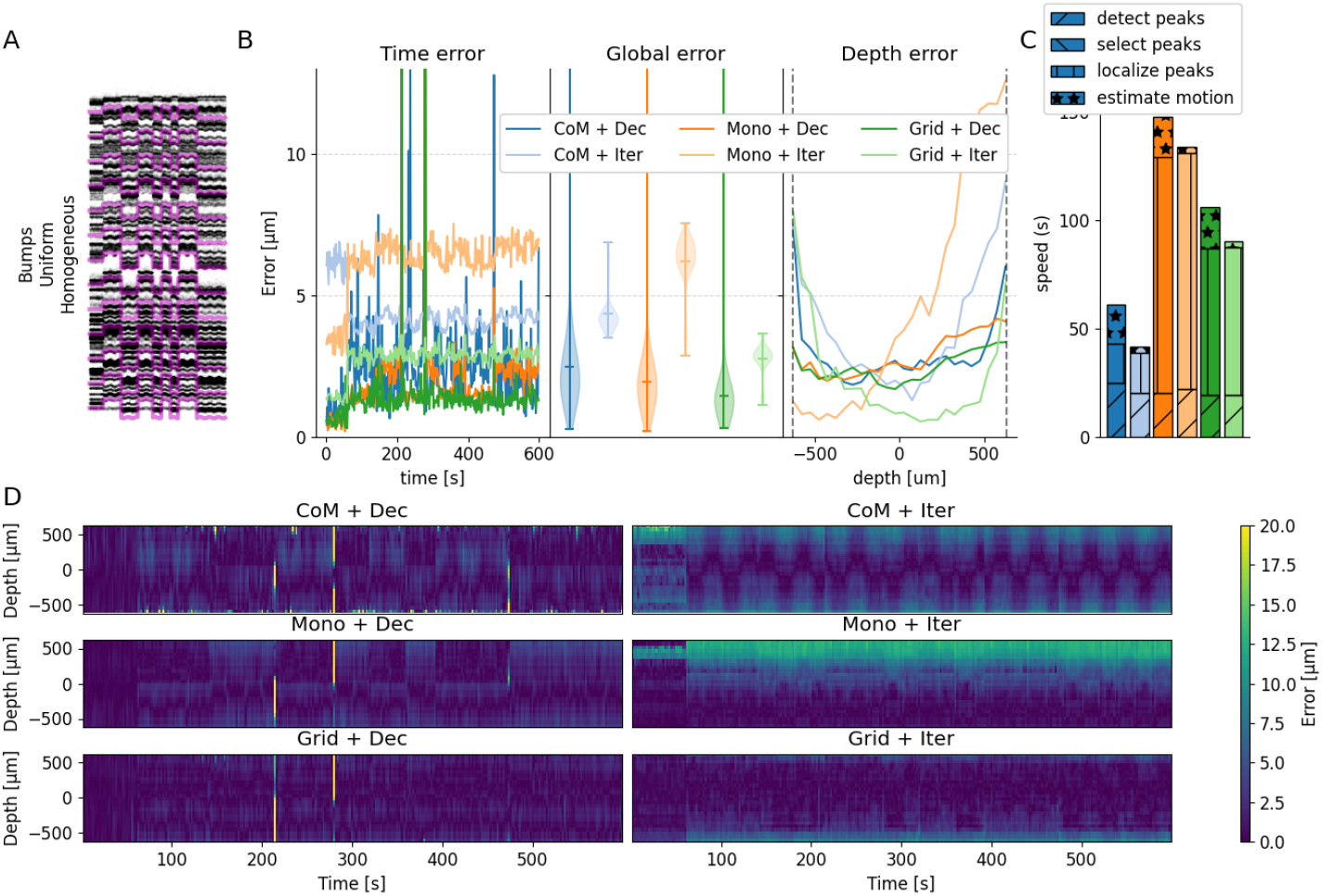
Motion estimation performance for the Bumps (non-rigid) drift case. **A)** Positional raster plot to show the movement of the cells, during the simulated drift. **B)** The errors with respect to the ground truth motion vector as function of the time (time error), averaged over time (global error), or as function of the depth (depth error) for several motion estimation pipelines. **C)** Run time of the different pipeline to estimate the motion (see legend below). **D)** Errors, as function of depth and time, for all the motion estimation methods considered.

To further investigate what is exactly happening during a complicated case, we decided to focus on the Bumps/Uniform/Homogeneous case in Figure 4. As one can see in Figure 4B, the errors made by the decentralized algorithm (either via *Mono* - orange - or *Grid* - green - localization) are lower, on average, compared to the ones made with the iterative template algorithm (see global errors). The method seems to be less sensitive to the depth (see Figure 4B, depth error), with lower errors on average. The computational cost of the motion estimation part is slightly higher (see Figure 4C), but most of the computational cost is due to the peak localization methods (the monopolar approximation is more accurate, but also slower as the results of an optimization problem). This is an important point not explored here, since we are computing the position for all peaks, but only in a 10-minute-long recording. For longer recordings, to limit the computational burden, sub-sampling the peaks prior to localization might be necessary (in the current implementation). The 4D panel shows the errors as function of both depth and time, for all motion estimation options. Large transient errors (yellow stripes) can be observed for the decentralized method (and also in one case for the iterative template method) and do not seem to be correlated with depths. Such large errors, although transient, can have a very negative impact at the spike sorting level, because during these time bins spikes are unlikely to be correctly recovered.

### Benchmarking motion interpolation

After quantifying the performance of motion estimation, we evaluated and compared different motion interpolation methods: snapping (Snap), inverse distance weighting (IDW), and kriging (Krig) (see Section “Motion interpolation” for details). Note that kriging is the method implemented in `Kilosort` [20].

As a representative example, we focused on one recording (Bumps/Uniform/Homogeneous) for this section. Figure 5A shows a portion of the raster maps (from 0 to 400 μm depth) for the *static* (grey) and drifting (red) recordings, and the interpolated recordings with kriging (green), IDW (orange), and snapping (blue). The raster maps are estimated from the interpolated traces after applying filtering, common median referencing, and z-scoring and using monopolar triangulation to estimate peak depths. To isolate the effects of interpolation, we used here the ground-truth motion vector as input for the various interpolation methods. Already from the raster maps, one can see that the kriging and IDW procedures seem to compensate well for the added motion, but in both cases some *wiggles* can be observed, indicating imperfections in the interpolation. In the ideal case, after interpolation one would expect straight bands similar to the *static* case. The snapping method, instead, is not capable to compensate for such large drifts, also due to the relatively low spatial density of the Neuropixels 1.0 configuration (with electrode distances of ∼20 μm). Higher electrode densities might improve the results of this method.

**Figure 5.**
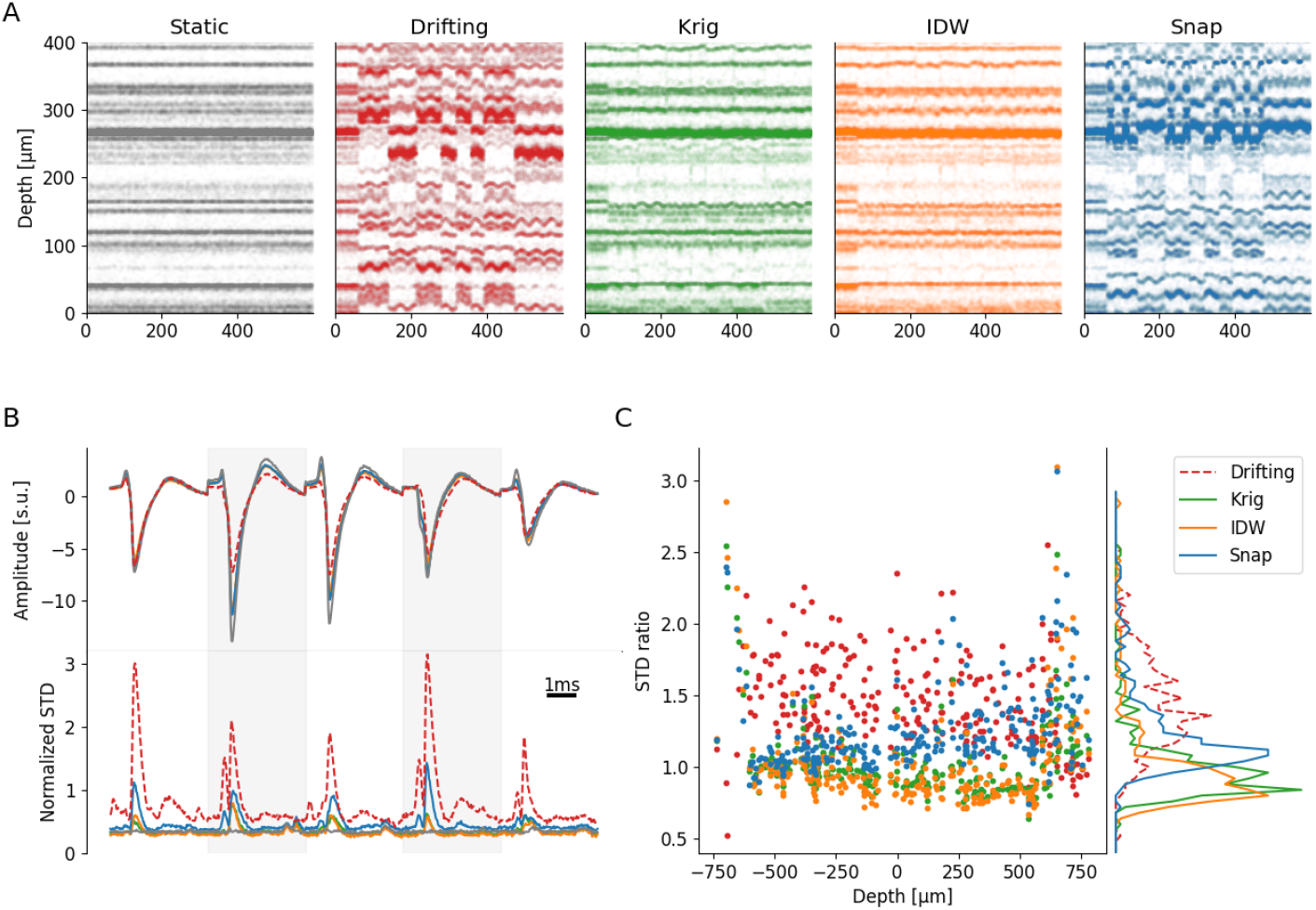
Quantification of the interpolation methods. **A)** Positional raster plots with a Bumps (non rigid) drift (uniform/homogeneous) for the static (grey), drifting (red), and interpolated traces via Kriging (green), IDW (orange) or Snap (blue), using the ground-truth motion vector. **B)** Top: Flattened template (on a subset of channels) for a single neuron in all the conditions (static, drifting, and the three interpolation methods), in units of the median absolute deviation (mad) of the noise on the considered channels. Bottom: Standard deviation over time, normalized by the template RMS, of all the individual spikes for this particular neuron, in the same conditions. **C** Ratios of waveform standard deviations compared to the static case for all the cells in the recording, in all conditions, and as function of the depth of the neurons. Right panel displays the overall STD ratio distributions. In this case, an STD ratio above 1 means an increase in waveforms dispersion with respect to the static case (and a decrease for ratios below 1).

To have a better insight on how interpolation affects spike waveforms, in Figure 5B we show the average template of one unit (top) and the temporal variance around the template (bottom) on the five channels with the largest amplitude. It is important to look at waveform variability since the main assumption of all spike sorting algorithms is that waveforms from a given neuron are reliable and stationary. A deviation from this assumption will therefore likely translate to worse spike sorting performances. For all interpolation methods, the interpolated template is smaller than the static counterpart (grey) and larger than the drifting recording (dashed red). This is probably due to the inevitable spatial smoothing that results from averaging the signal from different channels (for kriging and IDW). The temporal evolution of the standard deviation also shows an increase, specifically in correspondence of the template peaks. Nevertheless, the standard deviation is largely reduced with respect to the drifting case, indicating that motion interpolation *should* improve spike sorting results.

In Figure 5C, we show, for each neuron, the ratio of the waveform standard deviations with respect to the static waveforms, depending on the neuron depth (left) and the distribution of these ratios (right) for different cases (red - drifting, green - kriging, orange - IDW, blue - snapping). We remind that this view is an idealized situation in the drift correction pipeline, because here the exact ground-truth motion vector is provided to interpolate the traces. As expected, the waveform variability is systematically increased for all cells in the drifting case compared to static. For IDW and kriging, the overall ratio not only is reduced, but it is even less than one on average, probably due the spatial smoothing resulting from the interpolation procedures, which reduces the additive uncorrelated noise. Since the test recording has non-rigid drifts, we can also observe a slight correlation with depth, with smaller variabilities as the depth gets positive (and the drift amount reduces). A strong border effect at the borders of the probe is also visible, with STD ratios exploding irregardless of the methods. This is also expected, since all motion interpolation procedures, close to the probe borders, can only rely on partial information to interpolate the traces.

### Global impact at the spike sorting level

After evaluating the performance at the motion estimation and motion interpolation stage, we now investigate how a complete drift correction pipeline affects spike sorting results. From the previous section, we showed that IDW and kriging are the best interpolation options. For this analysis, we used kriging since it is also used by `Kilosort` 2.5 and it therefore allowed us to perform a direct comparison.

We ran spike sorting using `Kilosort` 2.5 on three different datasets – ZigZag, ZigZag (non rigid), Bumps – all with uniform depth distributions and homogeneous firing rates (Figure 2A, C, and E). For each recording, we ran four spike sorting options, and note that the cell distributions and spiking activity for all three recordings is the same, resulting in the exact same *static* recordings:

- **Static - no interpolation:** spike sorting on the *static* recording, turning off motion correction in `Kilosort` 2.5
- **Using GT:** spike sorting on the drifting recording, using the GT motion signals and kriging for motion interpolation in `SpikeInterface`
- **Using Mono+Dec:** spike sorting on the drifting recording, using the estimated motion with monopolar triangulation + decentralized inference for motion estimation and kriging for motion interpolation in `SpikeInterface`
- **Using KS2.5:** spike sorting on the drifting recording, letting `Kilosort` 2.5 preprocess the recording and correct for drifts (equivalent to grid localization + iterative template inference for motion estimation and kriging for motion interpolation)

Figure 6A-D-G shows the accuracy of the spike sorting for all GT units (sorted by accuracy). For all test cases, we observe a lower performance on the drift-corrected recordings with respect to the static ones (red lines). This drop is particularly dramatic in the complex case of Bumps (Figure 6G). Nevertheless, running motion correction using the Mono + Dec estimation in `SpikeInterface` (green lines) produces better results than using the `Kilosort` 2.5 correction procedure (purple lines) and it yields similar performance to using the ground-truth motion signals for interpolation (blue lines). In some cases, the accuracy using the estimated motion is even slightly better than the one using GT signals. This could be explained by the fact that the estimated motion is fully data driven, while the effect of the GT motion on the traces can also depend on the neuron location, morphology, and relative orientation with respect to the electrodes. Note that even in all cases, the accuracy loss is rather large, indicating that the interpolation step is degrading spike sorting outputs.

**Figure 6.**
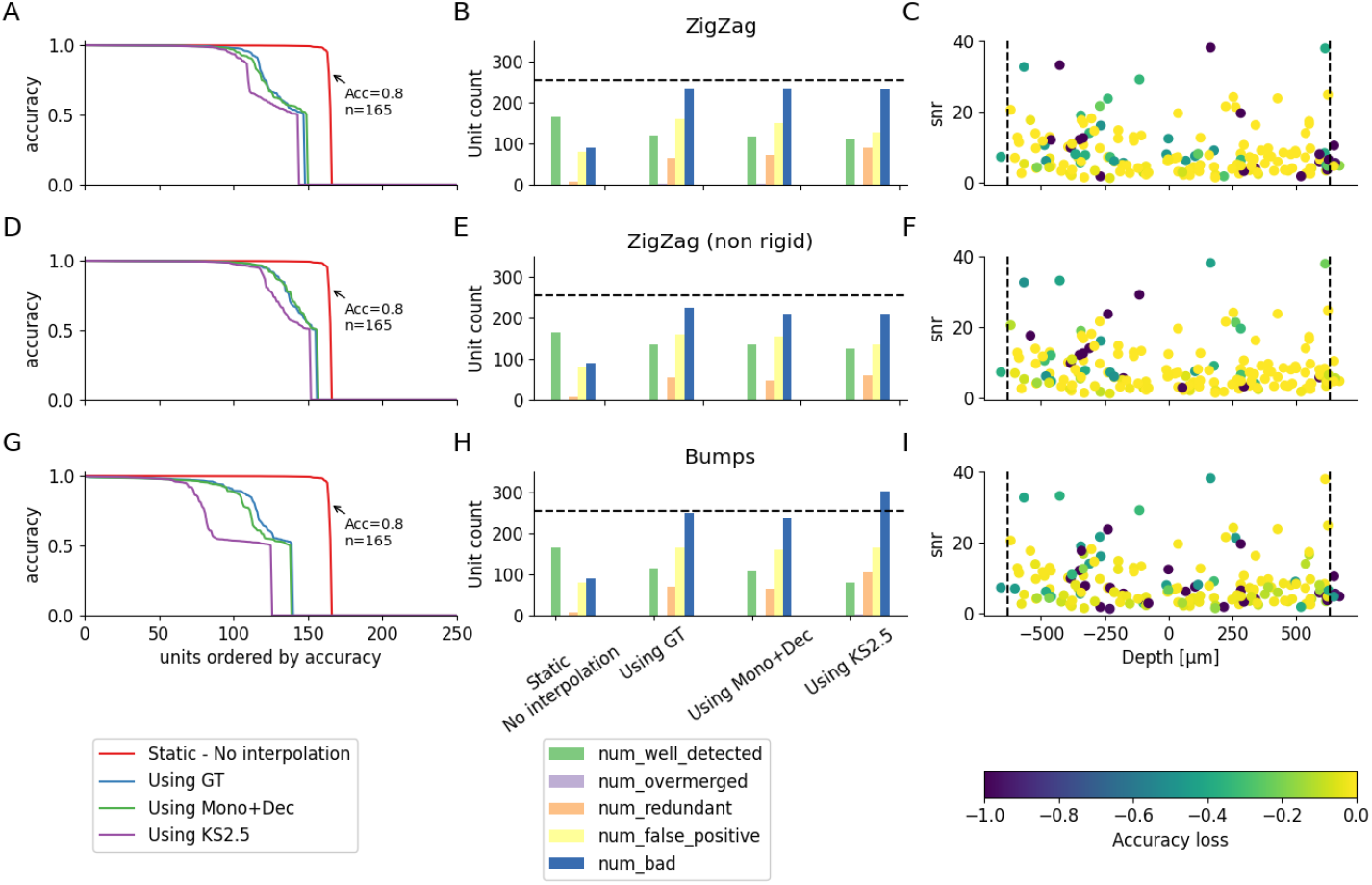
Quantification of the interpolation on sorting accuracy. **ADG)** Accuracy of all units (sorted by accuracy) in the simulated recordings when launching `Kilosort` 2.5 in various conditions for the Zigzag rigid case (**A**), the Zigzag (non-rigid case) (**D**) or the bumps (non-rigid) case (**G**). All recordings have uniform distributions and homogeneous firing rates. **BEH)** Number of well detected/overmerged/redundant/false positive and bad units found by `Kilosort`, with various drift correction methods, for the recordings explained in A, D, G. **CFI)** The loss in accuracy for the well detected neurons (accuracy higher than 0.8 - N=164) between the static case and the one using Mono+Dec estimation, as function of the depth of the neurons (x axis) and the signal-to-noise ratio (SNR - y axis).

When we look at the classification of sorted units (Figure 6B-E-H), the number of well detected units is 165 for the static recording, but it drops to: 118 for Mono+Dec and 110 for `Kilosort` on the ZigZag rigid recording; 136 for Mono+Dec and 124 for `Kilosort` on the ZigZag non rigid recording; and 107 for Mono+Dec and 79 for `Kilosort` on the Bumps recording. In addition, a much larger number of bad units is observed after drift correction, resulting from both false positive and redundant units. However, correcting with `SpikeInterface` also results in a reduced number of bad units with respect to `Kilosort` : for example, for the Bumps example, Mono+Dec finds 238 bad units and `Kilosort` finds 302.

Figure 6C-F-I we display the drop in accuracy for the well-detected units in the static recording (N=165) between the static and the *estimated Mono+Dec* cases depending on the unit depths (x axis) and signal-to-noise ratio (SNR - y axis). While a clear trend is not apparent, some units with large SNR seem to exhibit a large drop in accuracy. These drops in accuracy could partially be due to oversplits due to residual drifts, which would also explain the large number of redundant units. In order to assess this, we performed a merging procedure of the sorted results using the ground-truth information. In brief, for each ground-truth unit, all the sorted units with an accuracy greater or equal than 20% were merged into a single unit. Even after this merging step, there is still an observable drop in accuracy after interpolation compared to static (Figure S3A-D-G), but the number of well detected units is increased and redundant units are almost gone (Figure S3B-E-H) and the accuracy drops of high-SNR units are mostly recovered (Figure S3C-F-I). However, we want to highlight that this precise merging procedure is only possible for ground-truth data, and other strategies will need to be employed for experimental data, such as manual merging or alternative data-driven automated methods [17].

### Drift correction with SpikeInterface

In the previous sections we benchmarked the different steps involved in drift correction and their global impact on spike sorting outcomes. As mentioned in Section “A modular implementation of drift correction”, we implemented all the methods in `SpikeInterface` with a modular architecture. For example, here is a code snippet that shows how to estimate motion signals from a Neuropixels recording:

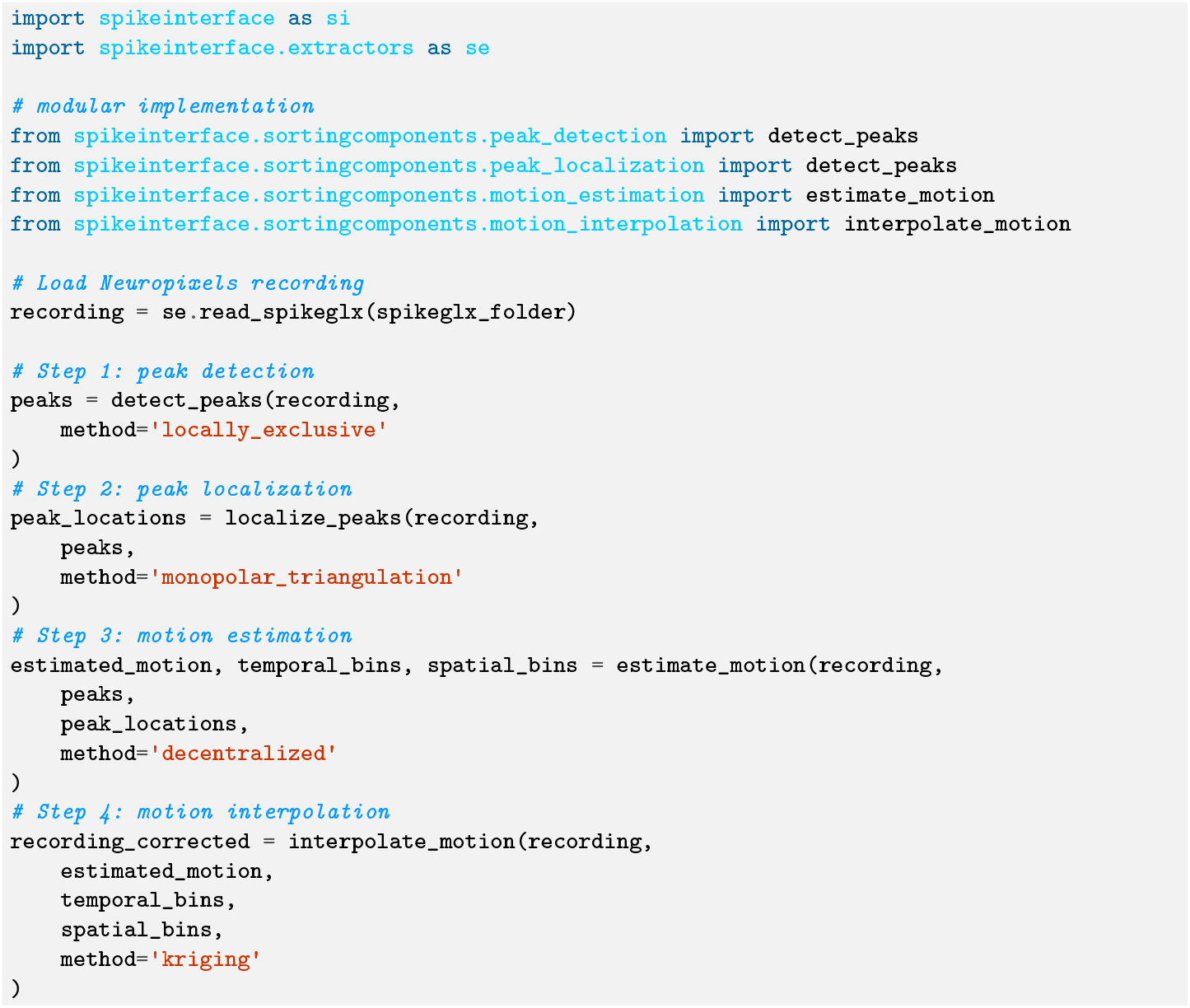

## Discussion

In this work, we have presented an exhaustive comparison of state-of-the-art methods to correct for motion in extracellular electrophysiology recordings. We generated a wide variety of simulated benchmark datasets using a Neuropixels 1.0 probe design with varying level of complexity, using Zigzag and/or *bumping* drift signals, rigid or non-rigid motion, and uniform or non-uniform depth distributions and firing rates. Knowing the ground-truth motion and spiking activity, we were able to finely evaluate the two different steps involved in drift correction, i.e. motion estimation and motion interpolation.

For motion estimation, we showed that the combination of monopolar triangulation (for peak localization) and decentralized registration generally outperforms all other options on the test datasets. Nevertheless, in its current implementation, localizing peaks with monopolar triangulation is more computationally expensive than other strategies (center-of-mass, grid interpolation), and the decentralized motion estimation method has a tendency to provide short but high spurious errors during some time bins. It is also important to notice that the accuracy in estimating motion signals is lower close to the borders of the probe, owing to the partial view of drifting activity with neurons that might move in and out of the *field of view* and thus be harder to track.

To evaluate motion interpolation, we compared the motion-corrected traces to a matching *static* simulation without added drift. We found that the *kriging* method (same interpolation method as `Kilosort` 2.5 [20]) achieves an equally good performance in reconstructing the static traces compared to inverse distance weighting scheme. Similarly to the drift estimation, the activity at the borders of the probe cannot be fully recovered/interpolated, because of missing information. Therefore, since both motion estimation and interpolation perform poorly at the borders of the probe, it would advisable, for recordings with apparent drift, to discard portions of the probe close to the borders from downstream analysis. The design of metrics and/or quantitative criteria to perform such a slicing of the probes should be the subject of further studies, and have already been implemented in `SpikeInterface`.

Finally, we evaluate the overall impact of the drift correction on spike sorting results. We found that applying the best motion estimation strategy (monopolar + decentralized registration) outperforms the `Kilosort` 2.5 implementation (grid interpolation + iterative template registration) and yields the same accuracy as using ground-truth motion signals for the motion interpolation step. Importantly, the reader should note that even applying the *best* drift correction dramatically reduces spike sorting accuracy with respect to a static recording. Electrophysiologists should therefore pay additional care in trying to minimize any major source of drifts, for example, by stabilizing the rig, lowering insertion speeds [10], and partially retracting the probe after insertion [9]. On the algorithmic side, re-interpolating the input traces to correct for drifts prior to spike sorting might not be the favorable solution moving forward. Instead, given the high accuracy of the motion estimation step, an alternative approach could use the motion signals *within* the spike sorting pipeline, for example by correcting waveform features before clustering or by using evolving templates over time in template matching.

In this work, we used only simulated datasets on Neuropixels 1.0-like probe designs as a benchmark. While clearly artificial data cannot fully substitute experimental ones, they provide a very controlled and customizable framework to test specific aspects of recordings, including drifts. A quantitative evaluation of drift correction methods on real data would in fact be very challenging due to the lack of ground truth. Even in the case of mechanical manipulation of the probe to inject drift [24], the relative movement of different layers of tissue with respect to the rigid probe movement would be hard to control. Furthermore, while here we focus on Neuropixels 1.0 probes, commercialized in 2019 and already widely used in neuroscience research, further benchmarks following a similar approach could target different probe layouts, such as Neuropixels 2.0 [25], Neuropixels Ultra (a prototype version with closely packed electrodes at 5 μm pitch), or other HD-MEA probe designs, such as the SiNAPS probe [1]. Furthermore, the presented simulation-based benchmark could be used in the *in silico* test and design of novel, more drift-resistant, probe layouts and geometries.

All motion estimation methods evaluated in this analysis use the detected peaks to construct activity histograms utilized to reconstruct drift signals. While these approaches are appropriate for recordings in rodents, they may fall short when dealing with recordings experiencing drifts at faster scales, such as the heart-beat and breathing modulations observed in recent recordings from humans [22] (similar drifts could be expected from non-human primates [26]). For drift at such fast timescales, local field potential (LFP) signals can be used instead of peak location estimates to build activity histograms [22, 29].

Finally, all the methods and drift correction options evaluated in this work are readily available to the electrophysiology community within the `SpikeInterface` package and can be immediately deployed with a few lines of code prior to any spike sorting process, as shown in Section “Drift correction with `SpikeInterface` “.

## Supplementary Figures

**Figure S1:**
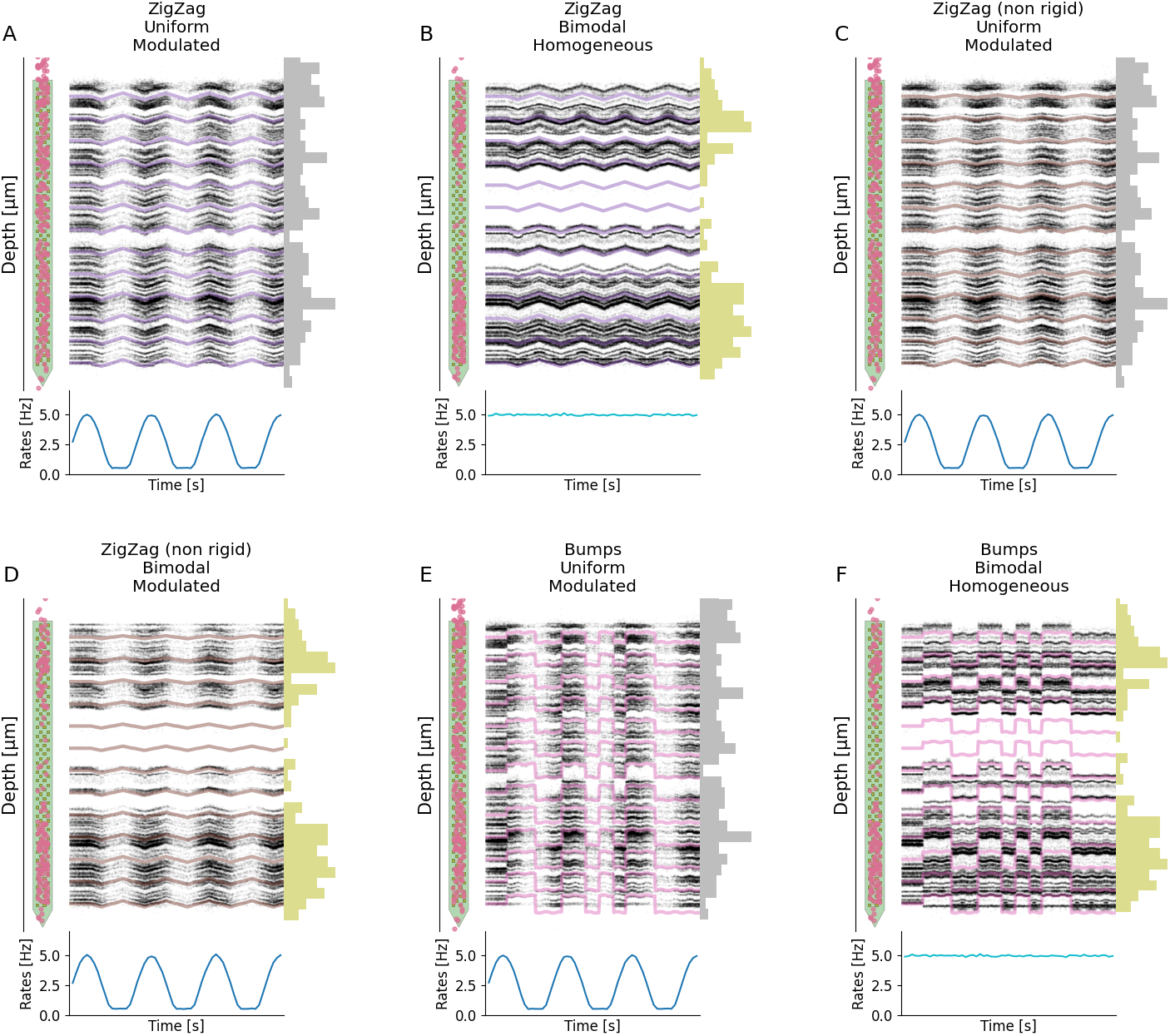
Other simulated drift recordings. For each panel, the top shows the layout of the probe with superimposed starting position of each cell (left), the positional raster plot with overlaid motion signals (center), and the depth distribution of the 256 neurons. The bottom part displays the firing rate modulation. **A)** Rigid ZigZag drift with uniform depth distribution and sine modulated firing rates. **B)** Rigid ZigZag drift with bimodal depth distribution and homogeneous firing rates. **C)** Non-rigid ZigZag with uniform depth distribution and modulated firing rates. **D)** Non-rigid ZigZag with bimodal depth distribution and modulated firing rates. **E)** Bumps with uniform depth distribution and modulated firing rates. **F)** Bumps with bimodal depth distribution and homogeneous firing rates.

**Figure S2:**
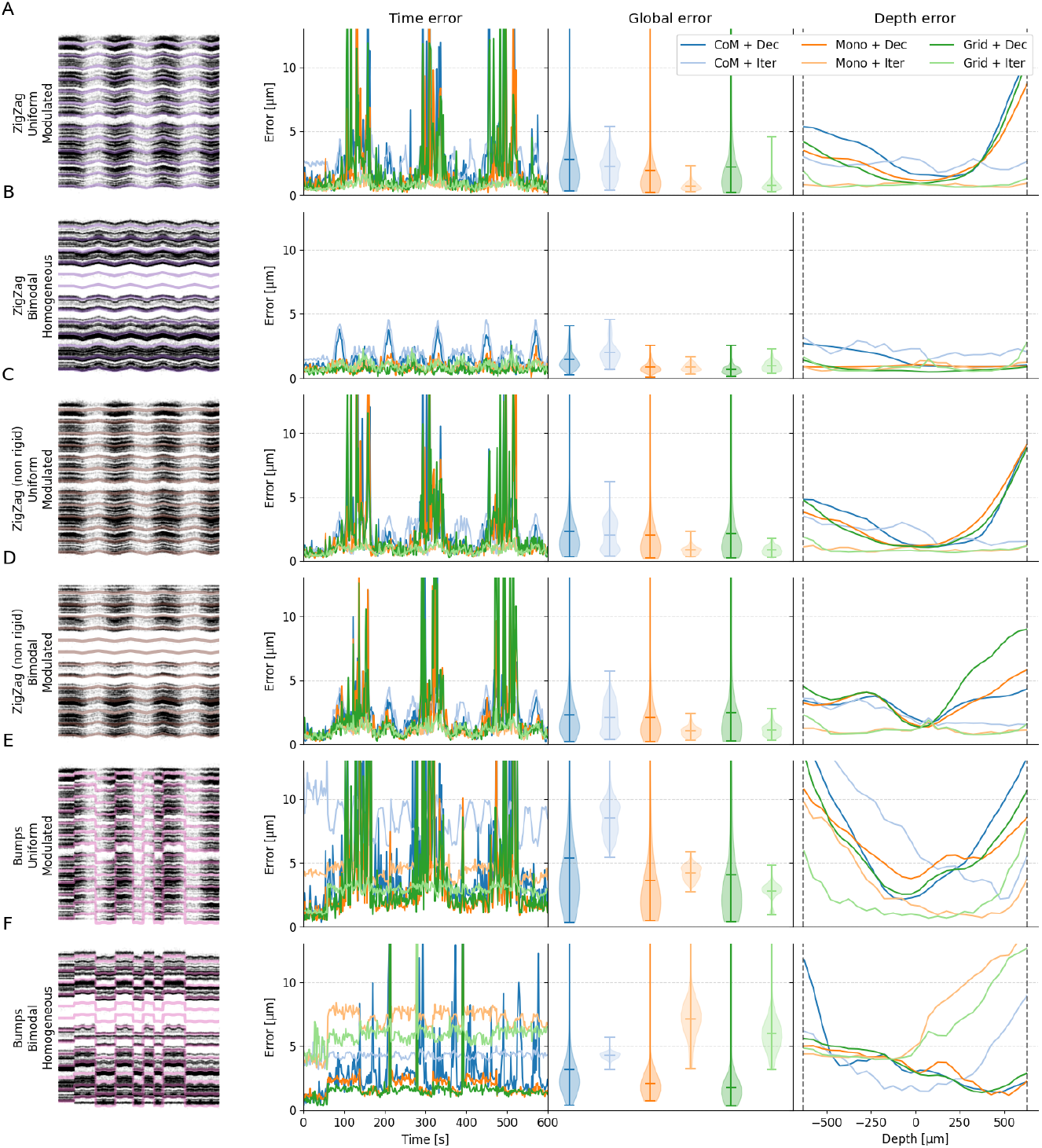
Performance in motion estimation for other drift scenarios. For various situations of the drifts (left column), errors (from left to right) as function of time, averaged, or as function of the depths and for various motion estimation pipelines. **A)** Zigzag (rigid), uniform positions, modulated rates **B)** Zigzag (rigid), bimodal positions, homogeneous rates **C)** Zigzag (non-rigid), uniform positions, modulated rates **D)** Zigzag (non-rigid), bimodal positions, modulated rates **E)** Bump (non-rigid), uniform positions, modulated rates **F)** Bump (non-rigid), bimodal positions, homogeneous rates

**Figure S3:**
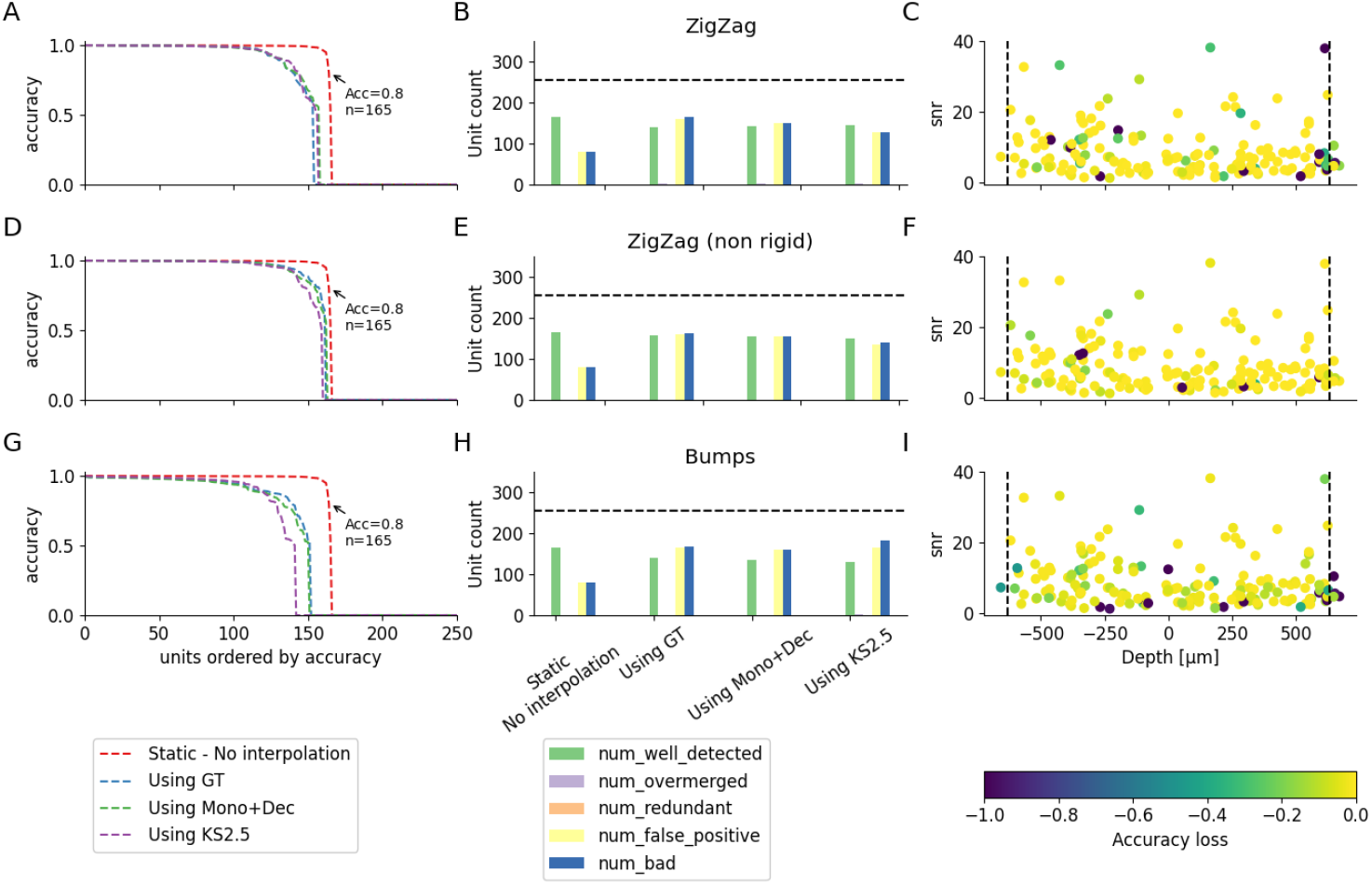
Quantification of the interpolation on sorting accuracy with quasi optimal merging. **ADG)** Accuracy of all units (sorted by accuracy) in the simulated recordings when launching `Kilosort` 2.5 in various conditions for the Zigzag rigid case (**A**), the Zigzag (non-rigid case) (**D**) or the bumps (non-rigid) case (**G**). All recordings have uniform distributions and homogeneous firing rates. **BEH)** Number of well detected/overmerged/redundant/false positive and bad units found by `Kilosort`, with various drift correction methods, for the recordings explained in A, D, G. **CFI)** The loss in accuracy for the well detected neurons (accuracy higher than 0.8 - N=164) between the static case and the one using Mono+Dec estimation, as function of the depth of the neurons (x axis) and the signal-to-noise ratio (SNR - y axis).

## Footnotes

1 the implementation was ported from pyKilosort - https://github.com/int-brain-lab/pykilosort/.

